# MGDb: An analyzed database and a genomic resource of mango (*Mangifera Indica* L.) cultivars for mango research

**DOI:** 10.1101/301358

**Authors:** Tayyaba Qamar-ul-Islam, M. Ahmed Khan, Rabia Faizan, Uzma Mahmood

## Abstract

Mango is one of the famous and fifth most important subtropical/tropical fruit crops worldwide with the production centered in India and South-East Asia. Recently, there has been a worldwide interest in mango genomics to produce tools for Marker Assisted Selection and trait association. There are no web-based analyzed genomic resources available for mango particularly. Hence a complete mango genomic resource was required for improvement in research and management of mango germplasm. In this project, we have done comparative transcriptome analysis of four mango cultivars i.e. cv. *Langra*, cv. *Zill*, cv. *Shelly* and cv. *Kent* from Pakistan, China, Israel, and Mexico respectively. The raw data is obtained through De-novo sequence assembly which generated 30,953-85,036 unigenes from RNA-Seq datasets of mango cultivars. The project is aimed to provide the scientific community and general public a mango genomic resource and allow the user to examine their data against our analyzed mango genome databases of four cultivars (cv. *Langra*, cv. *Zill*, cv. *Shelly* and cv. *Kent*). A mango web genomic resource MGdb, is based on 3-tier architecture, developed using Python, flat file database, and JavaScript. It contains the information of predicted genes of the whole genome, the unigenes annotated by homologous genes in other species, and GO (Gene Ontology) terms which provide a glimpse of the traits in which they are involved. This web genomic resource can be of immense use in the assessment of the research, development of the medicines, understanding genetics and provides useful bioinformatics solution for analysis of nucleotide sequence data. We report here world’s first web-based genomic resource particularly of mango for genetic improvement and management of mango genome.

## Introduction

As a member of the family Anacardiaceae, mango (*Mangifera Indica* Linn.) ranks second among 3 tropical fruit crops after banana due to its rich sensational taste, color, aroma and huge 4 economics significance (Litz RE, 2009; Srivastava S, 2016). According to Food and Agriculture Organization of the United Nations (FAO), India holds the 1^st^ position in mango production followed by China, whereas Pakistan and Mexico rank 5^th^ and 6^th^ position respectively (FAOSTAT-2014; www.faostat.fao.org). In the Ayurvedic and indigenous medical systems, it counts for an important herb for nearly over 4000 years (KA Shah, 2010). The substances in mango have high therapeutic potential. According to Ayurveda, mango tree comprises of different medicinal properties could be used to treat tumor, rabid dog, piles, toothache, diarrhea, cough, insomnia, hypertension, asthma, anemia, hemorrhage, dysentery, heat stroke, blisters, miscarriage, liver disorders, tetanus, wounds in the mouth and excessive urination. Phytomedicines should be sufficiently regulate based on this knowledge (Kalita, 2014).

Mango has a small genome size of 439 Mb (2n = 40) and a diploid fruit tree with 20 pairs of chromosomes (Arumuganathan, 1991). There are total 72 species of genus Mangifera from which most of them surviving in the rain forests of Malaysia and Indonesia (Nagendra K Singh, 2016). Mango has been widely cultivated in India and Southeast Asia for thousands of years. It has now grown throughout the world (tropical and sub-tropical) in 99 countries with a total fruit production of 34.3 million tons of fruit per annum (Galán Saúco, 2013). Asia is the largest producer of mangoes, 76% of world production comes from Asia with the second and third largest producers Americas (12%), and Africa (11.8%) (Galán Saúco, 2013).

Mango is a rich source of vitamins (vitamin A and vitamin C) and minerals and popular for its attractive color, texture and juicy flavor covers the total area of 2,297,000 ha in 2010 to 2011 (Hiwale1, 2015). The chemical analysis of mango pulp evinces that it has a relatively high content of calories (60 Kcal/100 g fresh weight) and is an important source of potassium, fiber, and vitamins (Marianna Lauricella, 2017). The nutritive value of mango is listed in (Table 1) reporting some data from the National Nutrient Database for Standard Reference (United States Department of Agriculture) (USDA, 2016).

**Table 1:** Nutritious Facts about Mangifera Indica L. (Mango) (Marianna Lauricella, 2017; USDA, 2016).

| **Mango Fruit Fresh, Nutrition Value per 100g** | **Nutrient Value** |
| --- | --- |
| **Energy** | 60 Kcal |
| **Carbohydrates** | 14.98 g |
| **Proteins** | 0.82 g |
| **Total Fat** | 0.38 g |
| **Cholesterol** | 0 mg |
| **Dietary Fiber** | 1.6 g |
| **Vitamins** |  |
| **Vitamin C** | 36.4 mg |
| **Vitamin A** | 1082 IU |
| **Vitamin E** | 1.12 mg |
| **Vitamin K** | 4.2 µg |
| **Niacin (vit B3)** | 669 µg |
| **Pantothenic acid (vit B5)** | 160 µg |
| **Pyridoxine (vit B6)** | 119 µg |
| **Riboflavin (vit B2)** | 38 µg |
| **Thiamin (vit B1)** | 28 µg |
| **Minerals** |  |
| **Sodium** | 1 mg |
| **Potassium** | 168 mg |
| **Calcium** | 11 mg |
| **Copper** | 110 µg |
| **Iron** | 160 µg |
| **Magnesium** | 10 mg |
| **Manganese** | 27 µg |
| **Zinc** | 90 µg |
| **Carotenoids** |  |
| **Carotene-β** | 445 µg |
| **Carotene-α** | 17 µg |

This project aims to build a successful response to the difficulties currently faced by fruit science agencies in meeting the need of gradually increasing fruit demands. The project intended to contribute to the development of a framework designed to support the analysis of the vast data for the availability of mango genomic information. It will also provide a platform for genetic information of different varieties of *Mangifera Indica* from different countries to make the estimated data available for the scientific community and general public.

The genomic data inside the mango genome database is derived from mango leaf transcriptome and chloroplast genome sequences estimated by different countries and reported in pieces of literature in 2014 (Azim MK, 2014). Mango fruit transcriptomes of cv. *Zill* (Wu HX, 2014), cv. *Shelly* (Luria N, 2014), cv. *Kent* (Dautt-Castro M, 2015), cv. Dashehari (Srivastava S, 2016) and more recently cv. *Keitt* (Tafolla-Arellano JC, 2017) have been reported from China, Israel, Mexico, India, and the USA respectively (Table 2).

**Table 2:** Statistics of unigenes data of mango cultivars (Waqasuddin Khan 2017).

| **RNA–Seq read**<br>**datasets** | **Total length of unigenes**<br>**(bases)** | **Total number of**<br>**unigenes** | **Length of largest**<br>**unigenes (bases)** | **No. of unigenes reported in the**<br>**respective paper** |
| --- | --- | --- | --- | --- |
| *cv. Langra* | 14,216,135 | 30,953 | 5,821 | 30,509 (Azim et al. 2014) |
| *cv. Zill* | 40,388,175 | 58,797 | 11,243 | 54,207 (Wu et al. 2014) |
| *cv. Shelly* | 49,681,799 | 57,544 | 8,723 | 57,544 (Luria et al. 2014) |
| *cv. Kent* | 58,412,829 | 85,036 | 8,305 | 80,969 (Dautt-Castro et al. 2015) |

Mango Genome Database comprises annotated data of species Mangifera *Indica* L. and analysis tools including the data of predicted genes, unigenes annotated in other species, related GO terms, Blast server, helpful bioinformatics tools with the user-friendly interface while ensuring the quality, maintainability, accessibility, and accuracy. The specific gene or annotated sequence data can be easily browsed or queried with various categories available in the gene search site. The genome sequence and the results generated after performing the queries can be easily accessed and downloaded. Mango Genome Database also provides analysis tools such as Blast server, Phylogeny, Plotting and Multiple sequence alignment. The aim of these tools is to allow the user to examine their data against our analyzed four mango cultivars (cv. *Langra*, cv. *Kent*, cv *Shelly* and cv. *Zill*) and to provide the best alignment and plotting tools with additional modification options with the facility of processing the results further. Here the analyzed genomic data of four cultivars of Mangifera *Indica* L. (cv. *Langra*, cv. *Kent*, cv *Shelly* and cv. *Zill*) will be available with the purpose of providing a separate unique platform of mango genome to the scientific community and general public where their data can be analyzed with different tools against the mango databases. The website will be the first website of particularly the mango genome and the first bioinformatics website created in Pakistan.

## Materials and Methods

### Retrieval of Mango unigenes Data

*Mangifera Indica* Linn. usually known as mango, is a species of the flowering plant in the sumac and poison ivy family Anacardiaceous. In this project four mango cultivars cv. *Langra* from Pakistan, cv. *Zill* from China, cv. *Kent* from Mexico and cv. *Shelly* from Israel transcriptomic sequences or the unigenes which are generated by processing the RNA-seq reads and transcriptome de novo assembly reported in 2014 (Azim MK, 2014) are retrieved, analyzed and presented in a database. We obtained four unigenes datasets corresponding to four mango cultivars from Jamil-ur-Rahman Center for Genome Research, Dr. Panjwani Center for Molecular Medicine and Drug Research, International Center for Chemical and Biological Sciences, University of Karachi. The statistics of the unigenes is shown in (Table 2).

### BLAST Analysis of Unigenes

The Blast Analysis or sequence comparison of unigenes of four mango cultivars were done against two databases: non–redundant (NR) protein sequence and Swiss–Prot. First, all unigenes of Pakistan dataset (cv. *Langra*) was aligned against non–redundant (NR) protein sequence database using BLASTX (E value cutoff ≤ 1e–3 and 10 hits per sequence) in Blast2GO java application (https://www.blast2go.com/start–blast2go–2–8) to retrieve common transcripts. Obtained results were then subjected to mapping and annotation for functional annotation and Gene Ontology (GO) assignments (Figure 1). This analysis was done only with cv. *Langra* from Pakistan for the Gene search module of the Website. Secondly, the four unigenes datasets are filtered for redundant sequences using CD-HIT (Cluster Database at High Identity with Tolerance) (Fu L, 2012) at 90% identity threshold. The resultant non-redundant unigenes were further analyzed for coding regions using TransDecoder (http://transdecoder.github.io/). Obtained coding sequences (CDS) were then subjected to homology search and BLASTed against Swiss–Prot databases by BLASTP with an E value cutoff ≤ 1e–3 to find the common and unique transcripts. These BLAST results are then used to modify the FASTA header of each sequence (unigene) of four assembled datasets (cv. *Langra*, cv. *Kent*, cv. *Shelly* and cv. *Zill*) (Figure 2). Following steps have been taken to modify the FASTA headers:

1. Extract the sequence, name and length columns from the original file.
2. Extract the description of the species with the name from the resultant file.
3. Merge the two files with the common column name.

**Figure 1:**
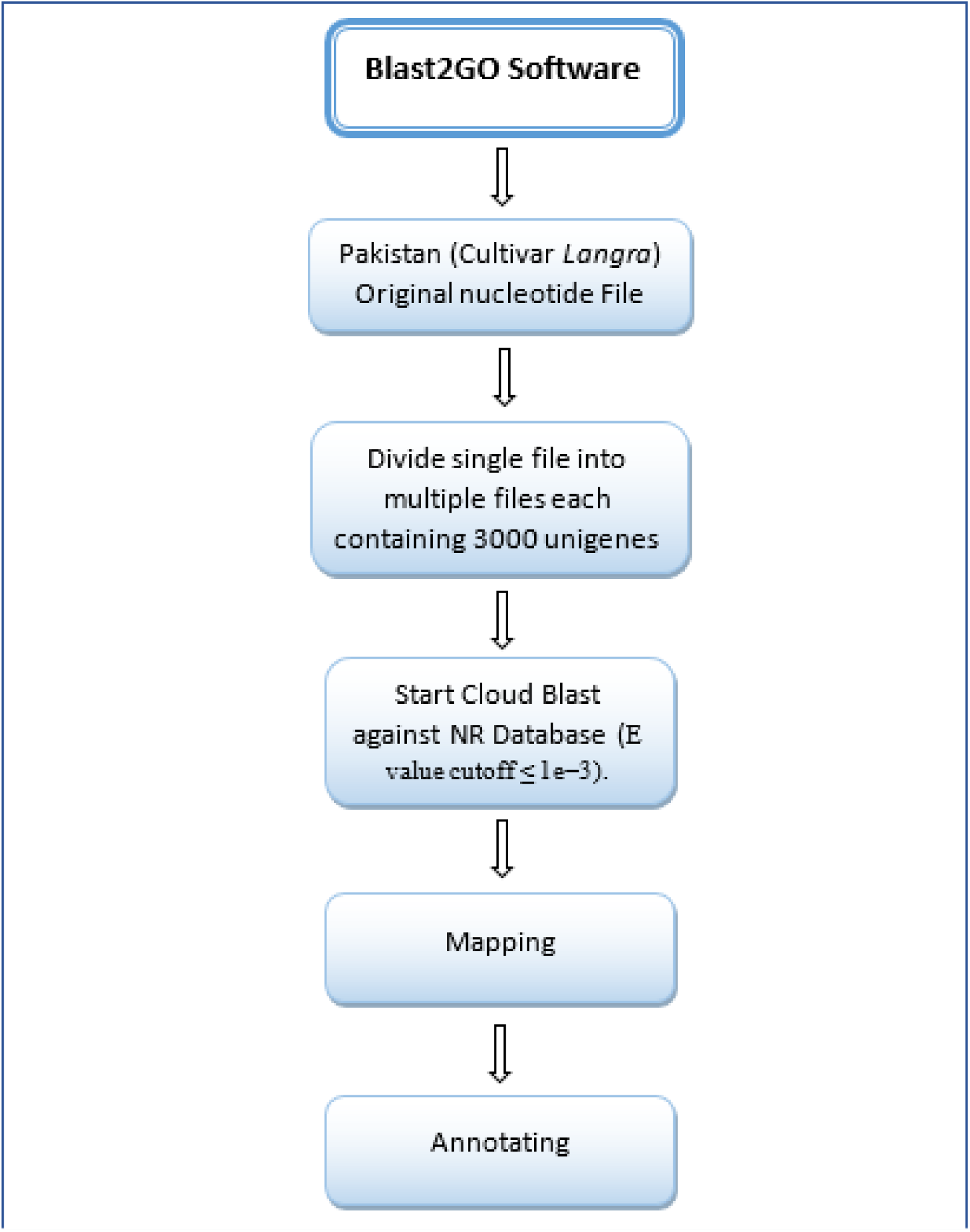
Flowchart schema of comparative analysis of Pakistan mango (cv. *Langra*) dataset.

**Figure 2:**
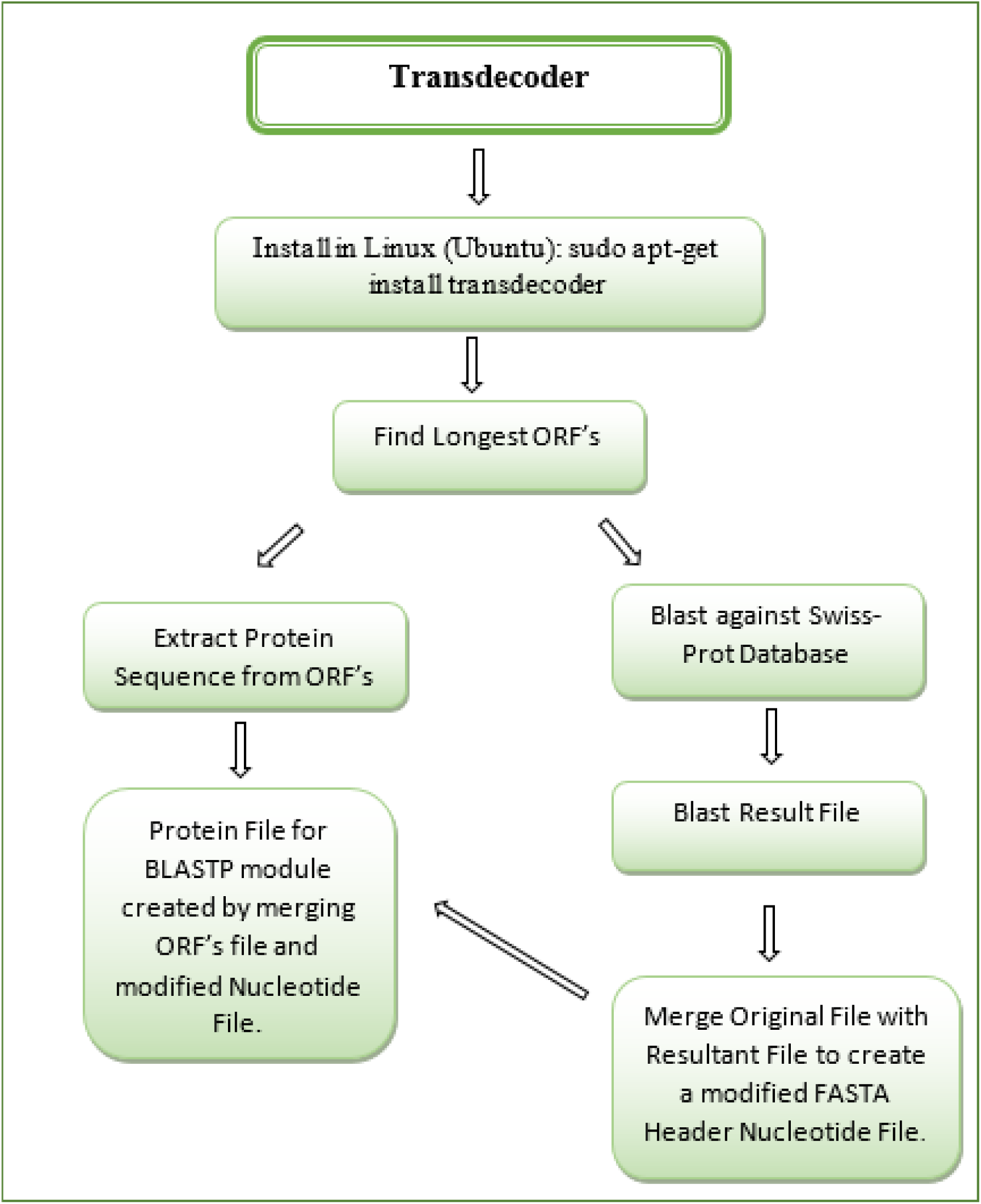
Flowchart schema of comparative analysis of four mango cultivars and formation of modified FASTA headers files.

### The Database Development

MGdb (Mango Genome database), a database with three-tier architecture was developed using Python, flat file database, and JavaScript. These three tiers are the client, middle and database tier. Multiple web pages and separate modules are developed for browsing and analysis. The user places the queries in client tier which sends the request to the middle tier that has a role in server-side scripting using python to fetch the data from a database (Iquebal et al., 2017). The database includes genomic data from different mango varieties (cv. *Langra*, cv. *Zill*, cv. *Kent* and cv. *Keit*).

### Construction of Analysis Tools

#### BLAST

The BLAST (Basic Local Alignment Search Tool) is an NCBI tool integrated into the website which allows the user to find the similarity of their entered sequences with our four analyzed unigenes datasets of mango cultivars. It employs a measure based on well-defined mutation scores. It uses the dynamic programming algorithm and directly approximates the obtained results for optimizing this measure. The method is faster than existing heuristic algorithm because it can detect weak but biologically significant sequence similarities and is one of the most used algorithms for alignment (Altschul SF, 1990). The search result will be shown in an easy-to-read table format and can be further expanded with a single click on the hit for related information and for an alignment view.

### Multiple Sequence Alignment

Multiple sequence alignment is a sequence alignment of three or more biological sequences, which could be DNA, RNA or protein. In Bioinformatics, sequence alignment is a standard technique for visualizing the relationships between residues in a collection of evolutionarily or structurally related sequences which are descended from a common ancestor (Waqasuddin Khan 2017). Here the different sequences of other species can be aligned with the mango cultivars unigenes dataset to identify their similarity with another genus. Multiple techniques of MSA (multiple sequence alignment) are used in this project which includes CLUSTAL W (http://www.genome.jp/tools-bin/clustalw), MUSCLE (https://www.ebi.ac.uk/Tools/msa/muscle) and EMBOSS (https://www.ebi.ac.uk/Tools/emboss/) programs of EMBL-EBI. The user can enter the sequences in FASTA format and select different options such as different output format, clustering method and maximum no of iterations etc. to get the desired results.

### Phylogenetic tree

The Phylogeny module of the website uses CLUSTAL W (http://www.genome.jp/tools-bin/clustalw) to do the alignment between multiple sequences. The tool is capable of handling some very difficult protein alignment problems. If the first alignment results are accurate or the data set consists of enough closely related sequences then CLUSTAL W will usually find an alignment that is very close to ideal (Thompson, 1994). This section of the website requires multiple sequences in FASTA format (either uploaded or pasted in a space given) and uses the ClustalW algorithm to find the ideal alignment. The results will be shown in a text and tree format with the facility to modify it with a different color of the clades and labeling and allow the user to extract more information from it. The results of the blast here can be further analyzed for the phylogenetic similarity and the results can be saved in different formats.

### Plotting

A plotting is a graphical technique and the pictorial representation of the data which evinces the relationship between two or more variables. After the analysis or alignment of the sequences, the user can plot the resultant file to see the graphical results. This section of the website includes the “compute sequence length” plot which uses the histogram tool to show the relationship between sequence length and base pairs. The user can use dot plot (simple and multiple sequence dot plot) to see the pictorial representation of the alignment of two or more sequences, similarity plot to find the similarity % of each residue in the sequence and GC % plot to find how much GC content is present in the uploaded gene sequence (Figure S2 and S3 in Additional File). The graphical results can be further amended to get the better view or the focused area.

## Results

The four mango cultivars from cv. *Langra*, cv. *Zill*, cv. *Shelly* and cv. *Kent* shows they have 30,953, 57,544, 58,797 and 85,036 unigenes respectively generated by transcriptomic de novo assembly resulted from Trinity (Grabherr MG, 2011; Waqasuddin Khan 2017). These unigenes were further characterized for functional annotations using BLAST (Jones P, 2014). BLAST homology search of cv. *Langra* showed 83% unigene sequences have 45-100% similarity with sequences in Nr database in which 59% sequences assigned with GO terms, 63% get mapped and remaining 17% are the novel sequences. The CDS analysis identified 14066 (~45%), 34893 (~59%), 58614 (~69%) and 35364 (~61%) protein coding sequences in cv. *Langra*, cv. *Zill*, cv. *Shelly* and cv. *Kent* unigenes datasets respectively. The homology search of these ORFs of four mango cultivars showed 57-100% similarity with sequences in Swiss-Prot database. The calculated probabilities for a variety of parameter choices by using the random model and scores are recorded in (Table 3) (Altschul SF, 1990).

**Table 3:** The probability in BLAST that the two random words of length (w) will have a score of minimum (T) (Altschul SF, 1990).

| W | T | Probability of a hit x 105 |
| --- | --- | --- |
| 3 | 11 | 235 |
|  | 12 | 147 |
|  | 13 | 83 |
|  | 14 | 48 |
|  | 15 | 26 |
|  | 16 | 14 |
|  | 17 | 7 |
|  | 18 | 4 |
| 4 | 13 | 127 |
|  | 14 | 78 |
|  | 15 | 47 |
|  | 16 | 28 |
|  | 17 | 16 |
|  | 18 | 9 |
|  | 19 | 5 |
|  | 20 | 3 |
|  | 16 | 28 |
|  | 17 | 16 |
|  | 18 | 9 |
|  | 19 | 5 |
|  | 20 | 3 |

## Discussion

Mango is one of the most important fruits of the tropical ecological region of the world, well known for its nutritive value, aroma, and taste (Iquebal et al., 2017). Transcriptomic sequences of different mango cultivars provided a wealth of data related to protein-coding sequences. This study and work resulted in an ‘analyzed web-based mango genomic resource’ from four mango cultivars grown in Pakistan, China, Israel and Mexico (Waqasuddin Khan 2017). Initially, we retrieved the mango genome assembled unigenes data of four different cultivars (cv. *Langra* (Pakistan), cv. *Zill* (China), cv. *Shelly* (Israel) and cv. *Kent* (Mexico)) from the international center for chemical and biological sciences located at the University of Karachi in Pakistan. These transcriptome sequences were obtained from RNA-Seq experiments using Illumina NGS technology (Waqasuddin Khan 2017). To analyze and annotate the data of mango transcriptome, total assembled unigenes from four different countries (cv. *Langra* (Pakistan), cv. *Zill* (China), cv. *Shelly* (Israel) and cv. *Kent* (Mexico)) were processed using three methods: comparative searching, mapping, and annotation.

### Comparison of assembled unigenes with Swiss-Prot and NR Databases

The total unigenes of 232329 from four different countries China, Pakistan, Mexico and Israel were first processed to find the ORF’s and then for the homology search in which the ORF results were BLASTed against Swiss-Prot database to find common transcripts which were then mapped and annotated to complete the process. The process was performed locally on Linux platform and the results are shown in (Figure 3 and Table 4) The dataset created by Pakistan contains the contigs sequences assembled into 30,952 unigenes. Mean size of unigenes was 536 bp with lengths in the range of 200 to >3,000 bp (Figure 4) (Azim MK, 2014). The process of comparing cv. *Langra* dataset of mango with Nr database starts by depositing the sequence file in Blast2GO java application (https://www.blast2go.com/start–blast2go–2–8) and blast it using BLASTX against Nr database. From 30953 unigenes 3000 sequences were get annotated from and the graph is created with the help of same blast2GO software (Figure 5 and S1 in Additional File).

**Figure 3:**
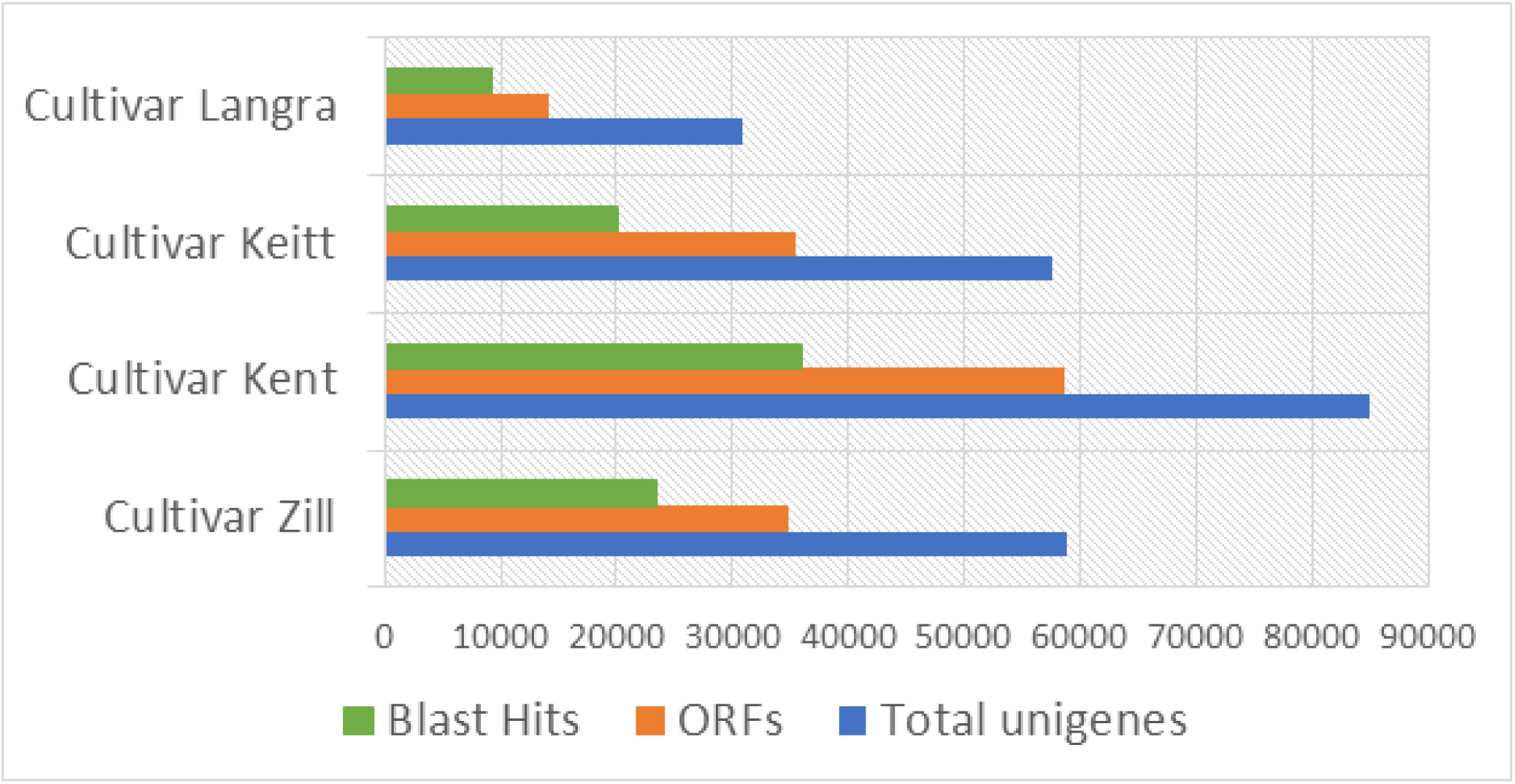
Results of blast search of four mango cultivars against Swiss-Prot database.

**Figure 4:**
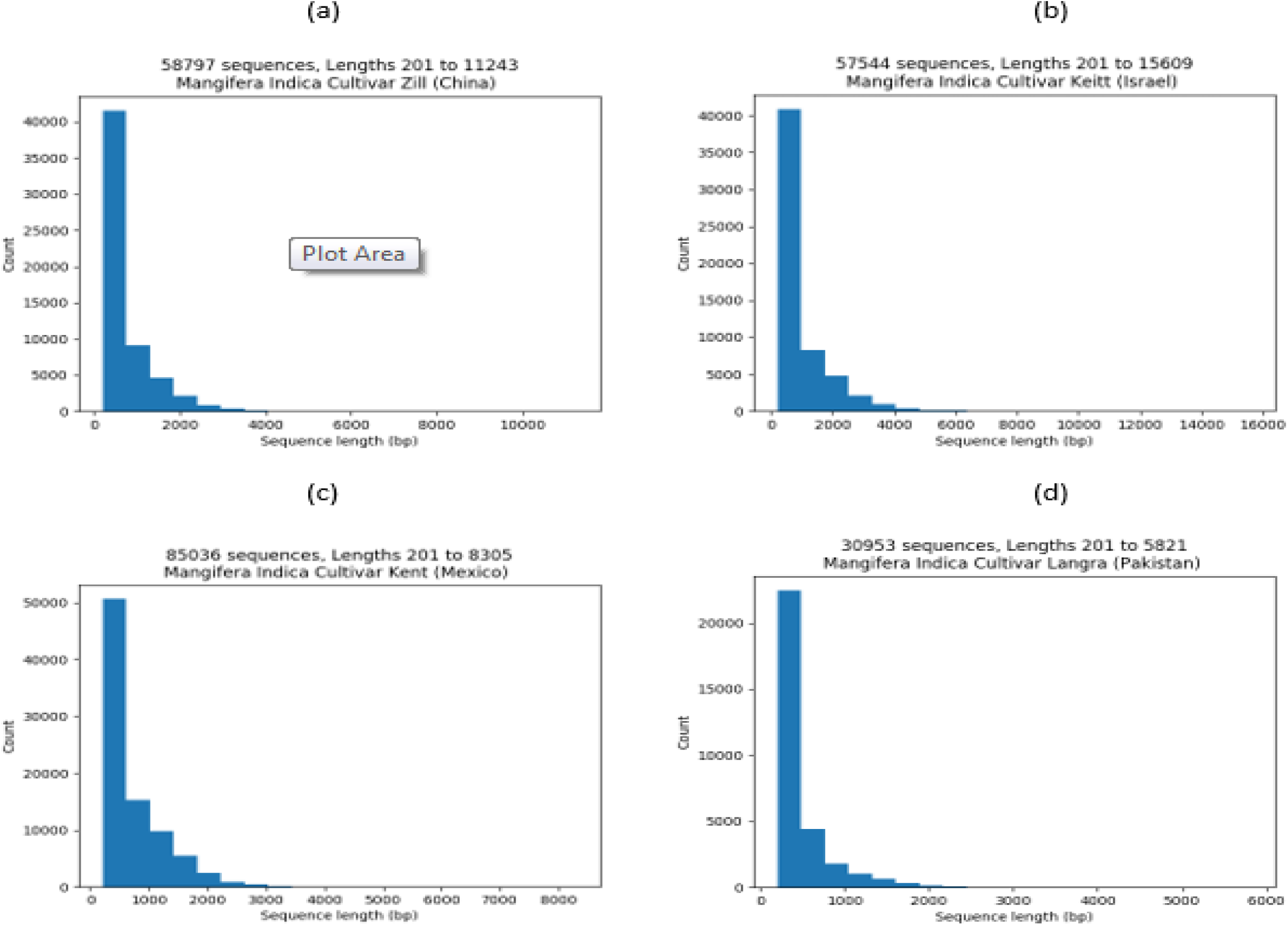
Length distribution of assembled mango unigene sequences of Mango (a) China (cv. *Zill*), (b) Israel (cv. *Shelly*), (c) Mexico (cv. *Kent*) and (d) Pakistan (cv. *Langra*).

**Figure 5:**
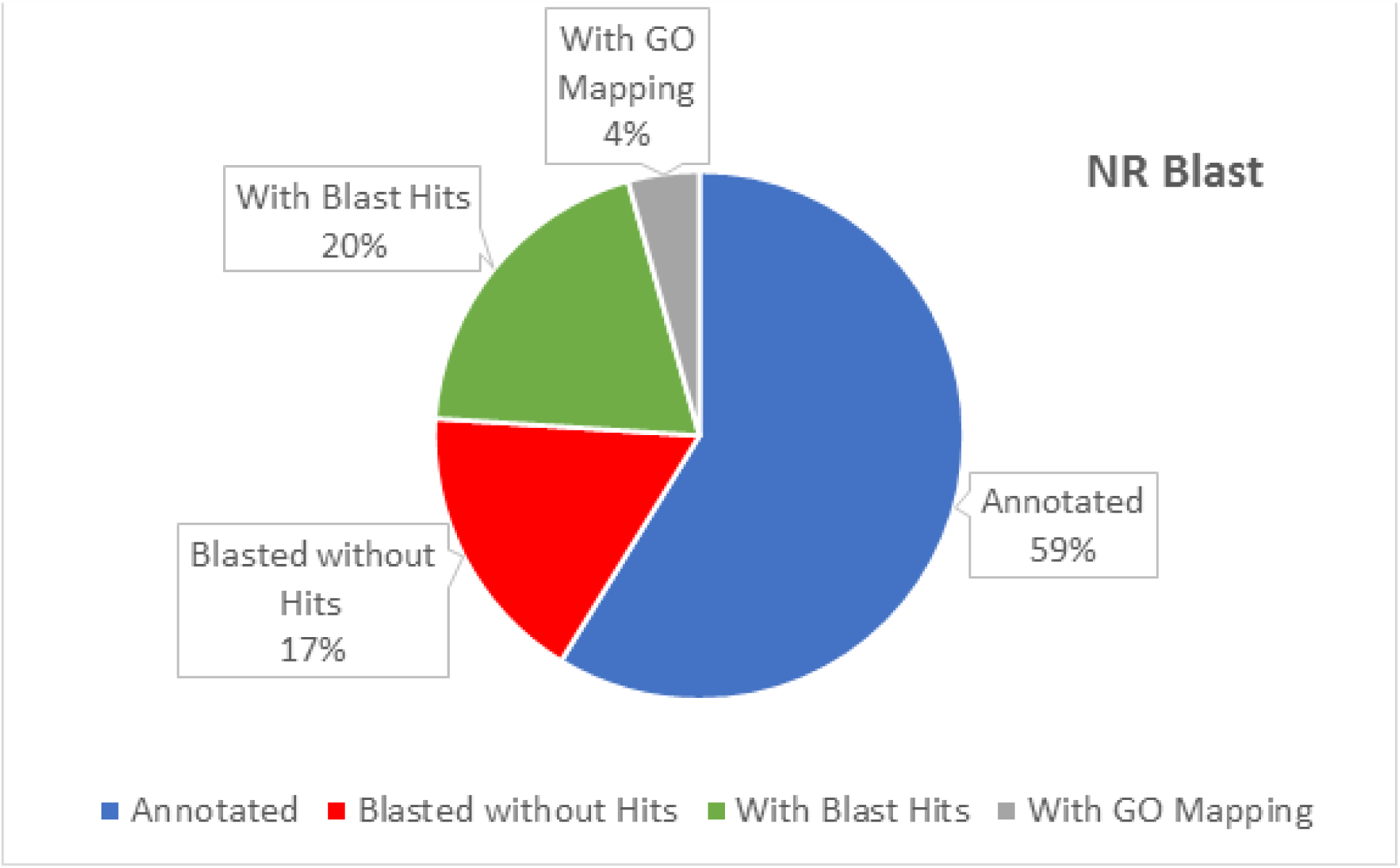
Blast search against NR database results represented in Pie Chart.

**Figure 7:**
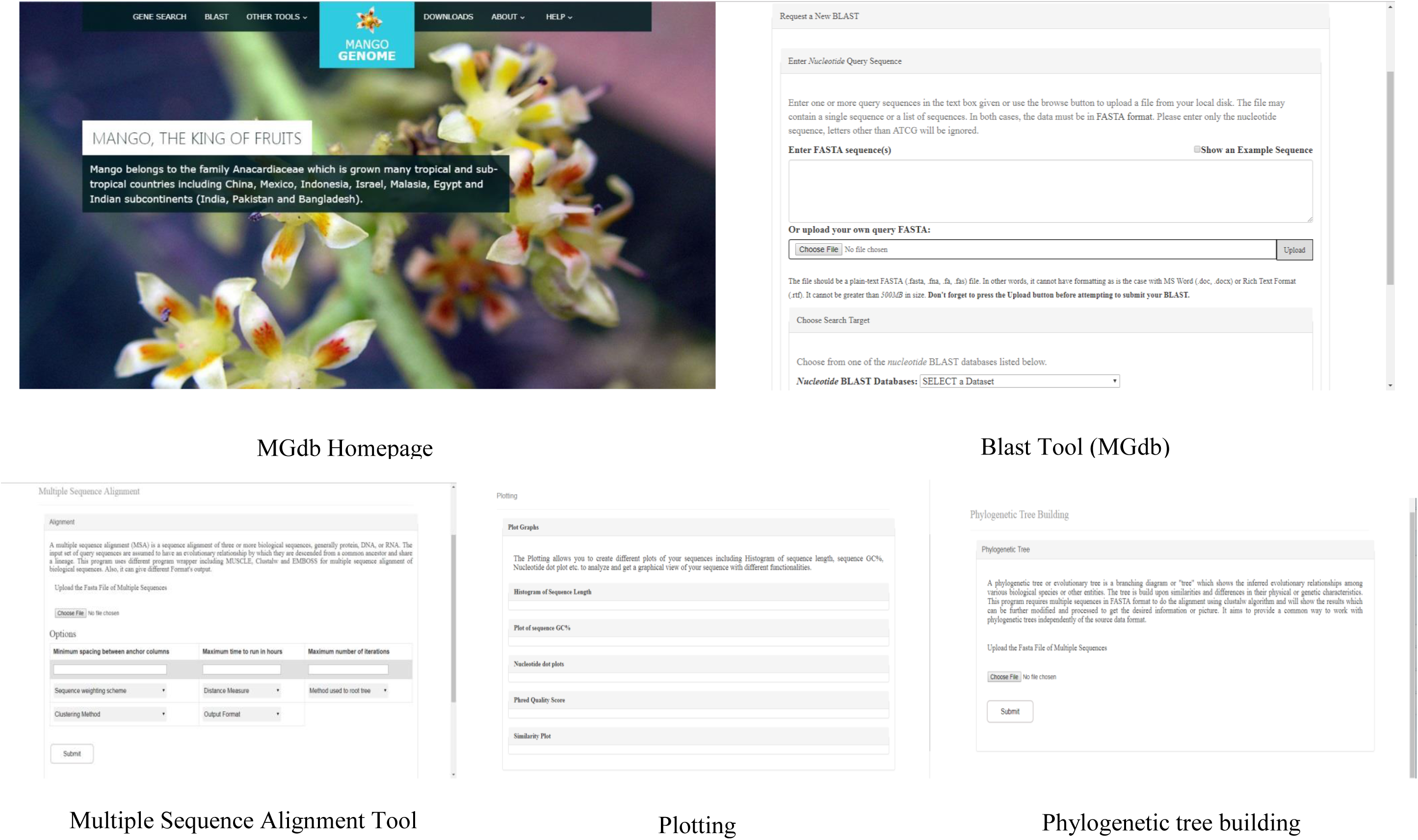

**Table 4:** Summary of mango unigene ORF’s and blast results from comparison with Swiss-Prot database.

| **Dataset** | **Total sequences** | **ORF's** | **Blast Hits** |
| --- | --- | --- | --- |
| Mangifera *Indica* L.<br>cultivar (Langra) | 30953 | 14066 | 9331 |
| Mangifera *Indica* L.<br>cultivar (Zill) | 58797 | 34893 | 34893 |
| Mangifera *Indica* L.<br>cultivar (Kent) | 85036 | 58614 | 36080 |
| Mangifera *Indica* L.<br>cultivar (Keitt) | 57544 | 35364 | 20064 |

BLASTX searching of mango unigene sequences against protein database (NR) was performed with E-value cut-off 10_–3_ and setting the no of hits to 15 from which the best one was considered for mapping and annotation. The result shows 24261 unigenes get the blast results out of total 30953 sequences which are then annotated to assign the GO terms to the Hits obtained through the BLAST search. These blast searches result in an XML file which is used to modify the FASTA headers of each dataset from different countries of *Mangifera Indica L* species (cv. *Langra* (Pakistan), cv. *Zill* (China), cv. *Shelly* (Israel) and cv. *Kent* (Mexico)). From 24261 BLASTed unigenes, 19531 unigenes were mapped from which 18188 get annotated. Functional characterization by Gene Ontology (GO) annotated an array of expressed genes in these mango varieties (Waqasuddin Khan 2017). Annotation is the process of selecting GO terms for the sequences from the GO pool assigned to one of the three biological domains (i.e. Biological, Cellular and Molecular functions) (Götz, 2007).

### Development of web-based mango genomic resource

Though a huge catalog of phenotypic information of mango and the related basic information is available in mango resources information system (www.mangifera.org), there are no web-based genomic resources available particularly of mango (Iquebal et al., 2017). The four datasets of mango cultivars (cv. *Langra* (Pakistan), cv. *Zill* (China), cv. *Shelly* (Israel) and cv. *Kent* (Mexico)) generated in the present study was analyzed and stored in a database (MGdb), which is the first portal available with genomic information of mango. The website is based on four sections: Analyzed sequences data, Blast, Gene Search, and other analysis bioinformatics tool. In the data section, the whole genome sequences, analysis results or the analyzed datasets of four mango cultivars of mango (cv. *Langra* (Pakistan), cv. *Zill* (China), cv. *Shelly* (Israel) and cv. *Kent* (Mexico)) can be accessed and downloaded in FASTA format. The user can use this entire data for the comparison of their own data of a different variety of mango or another species. The annotated sequence (assembled unigenes) data can be browsed through gene search page or queried using various categories in the search sites.

Blast (basic local alignment search tool) is a new approach to rapid sequence comparison which directly approximates alignments that optimize a measure of local similarity, the maximal segment pair (MSP) score (Altschul SF, 1990). The NCBI Blast tool used in a website allows the users to analyze their data against our analyzed mango genome datasets for finding multiple regions of similarity with certain DNA and protein sequences, motif searches and gene identification searches. The other analysis tools on the website include phylogeny, plotting, and multiple sequence alignment tool. Each tool has different input options, accepts FASTA and other mentioned format files for processing and provide the facility to transform the results according to the need (Figure 6).

Figure 6: Workflow of the MGDb genomic resource.

## Acknowledgment

We would like to express our sincere gratitude to Dr. Kamran Azim (project advisor) and Safina Abdul Razzak from Muhammad Ali Jinnah University for their invaluable advice, guidance and enormous patience throughout the development of the project.

We also acknowledge the support of Dr. Zaheer-ul-haq Qasmi from Dr. Panjwani Center for Molecular Medicine and Drug Research, International Center for Chemical and Biological Sciences, University of Karachi in the analysis of the data.

